# Identification of general patterns of sex-biased expression in *Daphnia*, a genus with environmental sex determination

**DOI:** 10.1101/269951

**Authors:** Cécile Molinier, Céline M.O. Reisser, Peter Fields, Adeline Ségard, Yan Galimov, Christoph R. Haag

**Author notes:** Equal contribution (Cécile Molinier, Céline M.O. Reisser). Corresponding author: Cécile Molinier, Centre d’Ecologie Fonctionnelle et Evolutive CEFE UMR 5175, 1919, route de Mende, 34293 Montpellier Cedex 5, France.

## Abstract

*Daphnia* reproduce by cyclic-parthenogenesis, where phases of asexual reproduction are intermitted by sexual production of diapause stages. This life cycle, together with environmental sex determination, allow the comparison of gene expression between genetically identical males and females. We investigated gene expression differences between males and females in four genotypes of *Daphnia magna* and compared the results with published data on sex-biased gene expression in two other *Daphnia* species, each representing one of the major phylogenetic clades within the genus. We found that 42% of all annotated genes showed sex-biased expression in *D. magna*. This proportion is similar both to estimates from other *Daphnia* species as well as from species with genetic sex determination, suggesting that sex-biased expression is not reduced under environmental sex determination. Among 7453 single copy, one-to-one orthologs in the three *Daphnia* species, 707 consistently showed sex-biased expression and 675 were biased in the same direction in all three species. Hence these genes represent a core-set of genes with consistent sex-differential expression in the genus. A functional analysis identified that several of them are involved in known sex determination pathways. Moreover, 75% were overexpressed in females rather than males, a pattern that appears to be a general feature of sex-biased gene expression in *Daphnia*.

**Short summary:** In some species with environmental sex determination, gene expression can be compared between genetically identical males and females. Here, we investigated sex-biased expression in one such species, *D. magna*, and compared it with data from two congeners. We found that all three species have a common set of 675 genes with consistent differential expression and with a strong bias towards overexpression in females rather than males. Moreover, the proportion of sex-biased genes in each of the three *Daphnia* species was similar to *Drosophila* species with genetic sex determination, suggesting that sex-biased expression is not necessarily reduced under environmental sex determination.

## Introduction

Patterns of gene expression often differ strongly between male and female individuals of the same species (Ellegren & Parsch 2007). In part, this difference is driven by genes on sex chromosomes, which show a particularly strong tendency for sex-biased (or sex-differential) expression (Ellegren & Parsch 2007; Bergero & Charlesworth 2009; Grath & Parsch 2016). However, sex-biased expression also occurs in many autosomal genes, and their products also fundamentally contribute to differences between male and female phenotypes (Ellegren & Parsch 2007; Grath & Parsch 2016; Wright *et al.* 2017). A particularly interesting case is that of species with environmental sex determination (ESD), where the same genotype may develop into a male or female, depending on environmental cues. Pure ESD species do not have sex chromosomes, and sex differentiation entirely relies on autosomal genes (Bull 1985). In GSD species, on the other hand, sex chromosomes contain a particularly high number of sex-biased genes (Mank 2009; Grath & Parsch 2016), and some of the genetic differences between sexes (e.g., sex-specific genomic regions or allelic differences) cause further, downstream expression differences, including for autosomal genes (Yang *et al.* 2006; Wijchers & Festenstein 2011). Both these observations suggest that species with ESD may have a lower number of genes with sex-biased expression compared to species with GSD. Alternatively, however, species with ESD may show a higher number of genes with sex-specific expression than species with GSD, because no genetic differences exist between sexes, and their entire sex-specific phenotypes are by differential gene expression (Grath & Parsch 2016, Mank 2017).

Among the species with ESD that have so far been investigated for sex-biased expression (Torres Maldonado *et al.* 2002; Shoemaker *et al.* 2007; Yatsu *et al.* 2016; Radhakrishnan *et al.* 2017), we find two species of the genus *Daphnia* (Colbourne *et al.* 2011; Huylmans *et al.* 2016). *Daphnia* reproduce by cyclical parthenogenesis: during the asexual phase of the life cycle, mothers clonally produce sons or daughters, but this asexual phase is intermitted by sexual reproduction, leading to the production of diapause eggs, which give rise to female hatchlings. Males and females are morphologically distinct (Scourfield & Harding 1966), and the sex of the clonally produced offspring is determined by environmental factors such as shortened photoperiod and/or increased population density (Roulin *et al.* 2013; Korpelainen 1990; Hobaek & Larsson 1990). Specifically, the production of males is induced by a hormone emitted by the mother in response to environmental conditions (Olmstead & Leblanc 2002). Moreover, male production can also be induced experimentally by adding the hormone analogue methyl farnesoate (MF) to the culture medium at a precise moment of the ovarian cycle corresponding to 48 to 72h after moulting (Olmstead & Leblanc 2002).

Since the publication of the *D. pulex* genome (Colbourne *et al.* 2011), the amount of genomic and transcriptomic resources for the genus has markedly increased (Routtu *et al.* 2014; Xu *et al.* 2015; Dukić *et al.* 2016; Orsini *et al.* 2016; Huylmans *et al.* 2016; Giraudo *et al.* 2017; Lynch *et al.* 2017; Spanier *et al.* 2017; Toyota *et al.* 2017; Ye et al. 2017; Herrmann *et al.* 2018). Previous studies have investigated sex-biased gene expression in two *Daphnia* species (Colbourne *et al.* 2011; Eads *et al.* 2007; Huylmans *et al.* 2016). The two species, *D. pulex* and *D. galeata* each belong to one of the two major phylogenetic groups within the subgenus *Daphnia* (Colbourne & Hebert 1997; Ishida, Kotov & Taylor 2006; Adamowicz, Petrusek & Colbourne 2009). These studies reported a high number of genes with sex-biased expression in both species and a preponderance of female-biased genes (i.e., genes overexpressed in females) compared to male-biased genes.

The genus *Daphnia* contains a second subgenus, Ctenodaphnia, which notably contains the species *D. magna.* This species is not only one of the major genomic model organisms of the genus (Miner *et al.* 2012, GenBank accession number: LRGB00000000.1), but also for studies on sex differentiation under ESD and other sex-related traits, including local adaptation in male production, evolution of genetic sex determination (which occurs in some genotypes of D. *magna*, not investigated here), and uniparental reproduction (Kato et al. 2011; Galimov, Walser & Haag 2011; Svendsen et al 2015; Reisser et al. 2016; Roulin *et al.* 2013). Here we present an analysis of sex-biased gene expression in *D. magna*, based on RNA-sequencing of males and females of four different genotypes (males and females of the same genotypes are members of the same clone). The four genotypes were used as biological replicates, that is, the expression of individual genes was classified as “sex-biased” only if the bias was consistent among all genotypes tested (i.e., if the overall pairwise test with four replicates was significant). While the primary aim was to study sex-biased gene expression in this important model organism, we also compare our results to those from the previous studies on *D. pulex* and *D. galeata*. The aims of this comparison were to identify general patterns of sex-biased gene expression in the genus, and to identify a core-set of genes with consistent sex-biased expression within the genus.

## Material and Methods

### Study design and origin of clones

We carried out RNA sequencing on adult males and females of *D. magna*, reared under standard culturing conditions (see below). We used four different genotypes which originated from a single population in Moscow, Russia (N55°45’48.65’, E37°34’54.00’’), as biological replicates. One library was prepared per genotype and sex, resulting in a total of eight libraries. Furthermore, each library consisted of eight technical (experimental) replicates, that is, eight clonal individuals of the same genotype and sex, pooled together into the library. Hence, a total of 64 individuals were raised for the experiment. Technical replicates were used to reduce variation due to small differences in environmental conditions (light, temperature, food, etc.) on gene expression. Such small environmental differences may be caused, for instance, by different positions of individuals within the culture tubes in the culture chamber.

### Laboratory culturing

Gravid parthenogenetic females were transferred individually to standard culturing conditions: a single individual in a 50mL Falcon tube containing 20 mL of artificial medium for *Daphnia* (Klüttgen *et al.* 1994), fed with 150 μL of algae solution (50 million of cells of *Scenedesmus* sp. per mL), and kept at 19°C under a 16:8 hour light-dark photoperiod. Each technical replicate was reared under these standard conditions during two pre-experimental clonal generations to remove potential maternal effects (Gorbi *et al.* 2011). To that end, one randomly selected offspring of the second clutch was transferred to a new tube to start the next clonal generation. Third-generation offspring were used for RNA sequencing.

Third generation males were produced by placing second generation females in a medium containing 400 μM of methyl farnesoate (“MF”, Echelon Biosciences) just prior the production of their first clutch, as determined by well-swollen ovaries (Olmstead & Leblanc 2002). This ensured that the sex of their second clutch offspring was determined in the presence of MF. Otherwise, males were treated in the same way as described for the females. In particular, the newborn males were transferred back to standard medium, just as the third-generation females, and, throughout, culture media were exchanged daily for both males and females.

### Sampling

Just before the third-generation females released their first clutch, all individuals (males and females) were transferred individually to a 1.5 mL well on a culture plate, where they were kept for about three days before being sampled. Since RNA was extracted from whole individuals, no food was added during the last 12 h before sampling in order to minimize algal RNA contamination (most of which will be digested and hence degraded after 12 h). The period without food was kept relatively short to minimize starvation-dependent gene regulation.

To remove as much culture medium as possible, the individuals were blotted with absorbing paper (previously sterilized with UV radiation for 30 minutes), and then transferred to a 1.5 mL tube that was directly immersed in liquid nitrogen. Directly after the flash-freezing, three volumes of RNAlater ICE solution were added to preserve RNA, and samples were placed at −80°C.

### RNA extraction, library preparation, and RNA-sequencing

The eight technical replicates (individuals of the same genotype and sex) were pooled, resulting in two samples (one male, one female) per biological replicate. Total RNA extraction and purification was carried out for each of the eight samples following the protocol of the *Daphnia* genomic consortium (DGC; DGC, Indiana University October 11, 2007), using Trizol Reagents and Qiagen RNEasy Mini Kit. The extracted and purified RNA samples were transferred to −80°C and shipped on dry ice to the BSSE Genomic Sequencing Facilities, University of Basel, Switzerland.

Two lanes of cDNA corresponding to the eight biological replicates were constructed by the Department of Biosystems Science and Engineering (D-*BSSE*). The eight samples were individually labelled using TruSeq preparation kits. Each library (2 lanes) was sequenced on an Illumina NextSeq 500 sequencer with 76 cycles with the stranded paired-end protocol.

### Quality control and filtering

Read quality was assessed with FastQC v.0.10.1 (http://www.bioinformatics.babraham.ac.uk/projects/fastqc). Paired-end sequences were subjected to adapter trimming and quality filtering using trimmomatic v.0.36 (Bolger, Lohse & Usadel 2014). After trimming of adapter sequences, terminal bases with a quality score below three were removed from both ends of each read. Then, using the sliding window function and again moving in from both sides, further 4 bp-fragments were removed while their average quality score was below 15.

### Mapping and counting

Filtered reads were mapped to the publicly available *D. magna* genome assembly (NCBI database; Assembly name: daphmag2.4; GenBank assembly accession: GCA_001632505.1, Bioprojects accession: PRJNA298946), consisting of 28’801 scaffolds, 38’559 contigs and a total sequence length of 129’543’483 bp, as well as a genome annotation with 26’646 genes, using the RNA-Seq aligner STAR (Dobin et al. 2013) using default settings. The raw counts (number of mapped reads per transcript per sample) were obtained with the software featureCounts (Liao, Smyth & Shi 2014), a fast tool for counting mapped paired-end reads. Counts were summarized at the gene level using the annotation file.

### Differential gene expression

The analysis of differential gene expression was carried out with DESeq2 (version 1.10.1, Love, Huber & Anders 2014) implemented in R (R Core Team 2017). Raw read counts were used as input data, and the subsequent analyses used the normalizations of read counts as performed by DESeq2, which is currently considered best practice for the analysis of RNA-sequencing data (Conesa *et al.* 2016; Schurch *et al.* 2016). The males vs. females comparison was carried out with a two-factor design taking into account clone identity and sex. All *p*-values were adjusted for multiple testing with the Benjamini-Hochberg method, as implemented in DESeq2. Genes were considered differentially expressed (DE) if they had an adjusted *p*-value < 0.05 (False discovery rate, FDR = 5 %). The degree of sex bias was determined by the fold-change (abbreviated FC) difference between the treatments. DE genes were classified into five groups: <2-fold, >2-fold, 2- to 5-fold, 5- to 10-fold and >10-fold difference in expression (absolute changes rather than log-transformed changes). To obtain a broader overview of the expression profiles of the significantly DE genes, we performed a hierarchical clustering of the sex-DE genes, as implemented in DESeq2. The normalization in DESeq2 does not account for transcript length, hence it is possible that some differential exon usage (that could ultimately result in the existence of different isoforms) could be mistakenly interpreted as differential expression. However, because different transcripts of most genes differ only weakly to moderately in size (Chern *et al.* 2006), normalization by transcript length (which has been criticizes for other reasons, Dillies *et al.* 2013) would not strongly affect the inferred levels of fold-change in expression levels. Therefore, inferences of differential expression should typically be robust, at least in the class of genes with a greater than two-fold change.

### Comparison of sex-biased gene expression with *D. galeata* and *D. pulex*

Protein sequences of *D. magna* (v2.4 GenBank: LRGB00000000), *D. pulex* (version JGI060905: http://wfleabase.org/release1/current_release/fasta/) and *D. galeata* (http://server7.wfleabase.org/genome/Daphnia_galeata/) reference genomes were used as input in the software OrthoFinder (Emms & Kelly 2015), a fast method for inferring orthologous groups from protein sequences with enhanced accuracy. These correspond to 26’646, 30’940, and 33’555 protein sequences, respectively. We also used another software, OrthoMCL (Li, Stoeckert & Roos 2003) for inferring orthologs. Because the results were qualitatively and quantitatively similar, we present only the results of OrthoFinder here. For further analysis, we retained only those genes that were identified by OrthoFinder as single copy, one-to-one orthologs in all three species. We then compared this list with our list of sex-DE genes (adjusted *p* < 0.05), as well as the lists of sex-DE genes in *D. galeata* and *D. pulex* (Colbourne *et al.* 2011; Huylmans *et al.* 2016). The R package VennDiagram (Chen & Boutros 2011) was used for visualization of sex-DE genes in the three species.

We then focused on the core subset of 675 orthologous genes that were found to be consistently sex-DE in all three species and used BLAST2GO (version 4.1.9, (Conesa *et al.* 2016) to perform a functional annotation. The protein sequences of the *D. magna* genes in question were annotated using the NCBI nr database, allowing for 20 output alignments per query sequence with an e-value threshold of 0.001. The mapping and annotation steps implemented in BLAST2GO were run with default settings. Additionally, InterPro IDs from InterProScan were merged to the annotation to improve accuracy. Graphical representation of GO categories belonging to the 675 core-genes was obtained using the R package metacoder (Foster, Sharpton & Grünwald 2017). The BLAST2GO output files were searched for the terms “sex determination” (GO accession number 0007530), “sex differentiation” (0007548), “male sex differentiation” (0046661), “female gonad development” (0008585), “male gonad development” (0008584), “female sex determination” (0030237), “male sex determination” (0030238), and “female sex differentiation” (0046660). In addition, a universal list of genes involved in sex determination and sex differentiation pathways was established by searching the entire Genbank database for genes associated with the terms “sex determination/differentiation”. After removal of redundancies, we obtained a list of 541 genes (hereafter referred to as the “NCBI list of genes”). We compared the annotations of our 675 core-genes (as determined by the BLAST2GO analysis) with this list to identify any shared genes. Finally, we performed an enrichment analysis of GO terms for the 675 core genes with consistent sex-biased expression in all three species, taking the 26’646 genes of *D. magna* as the reference GO composition. This was done using the GOatools Python script (https://github.com/tanghaibao/goatools), which performs Fisher’s exact tests for differences in frequencies of GO-terms between the two lists (with Bonferroni correction). Enriched GO categories were summarized by a reduction of the complexity and level of GO terms (medium; allowed similarity=0.7). UniprotKB was used to determine Gene Ontology Biological Process and Molecular Functions that were over-represented among genes DE between sexes and visualized with REVIGO (Supek *et al.* 2011).

## Results

### Data quality

The RNA sequencing of the eight libraries resulted in a total of 1.59 billion raw reads. An average of 99.01 % of raw reads passed the quality control. After end-trimming, an average of 93.03 % aligned to the reference genome, resulting in an average of 81 million aligned reads per library, which constitutes a robust data basis for differential gene expression analyses. **Table S1** shows the percentages of reads retained at each step in each of the samples.

### Sex-biased gene expression in *Daphnia magna*

We found a high number of genes that were DE between males and females with a total of 11’197 out of 26’646 genes being DE (adjusted *p* < 0.05), of which 8384 genes showed at least a two-fold change (**Table 1**). The strong sexually dimorphic expression patterns can be visualized in the expression heatmap of the 8384 DE genes with more than 2-fold expression difference between the sexes (**Fig. 1**). Overall, a slight, but significant (*p* < 0.0001) majority of those genes were male-biased rather than female-biased (**Table 1**). This male-bias was found for all categories, except for the genes with a weak (< 2-fold) sex bias (**Table 1**). The list of all sex-biased genes can be found in supplemental data (**Table S2**).

**Figure 1:**
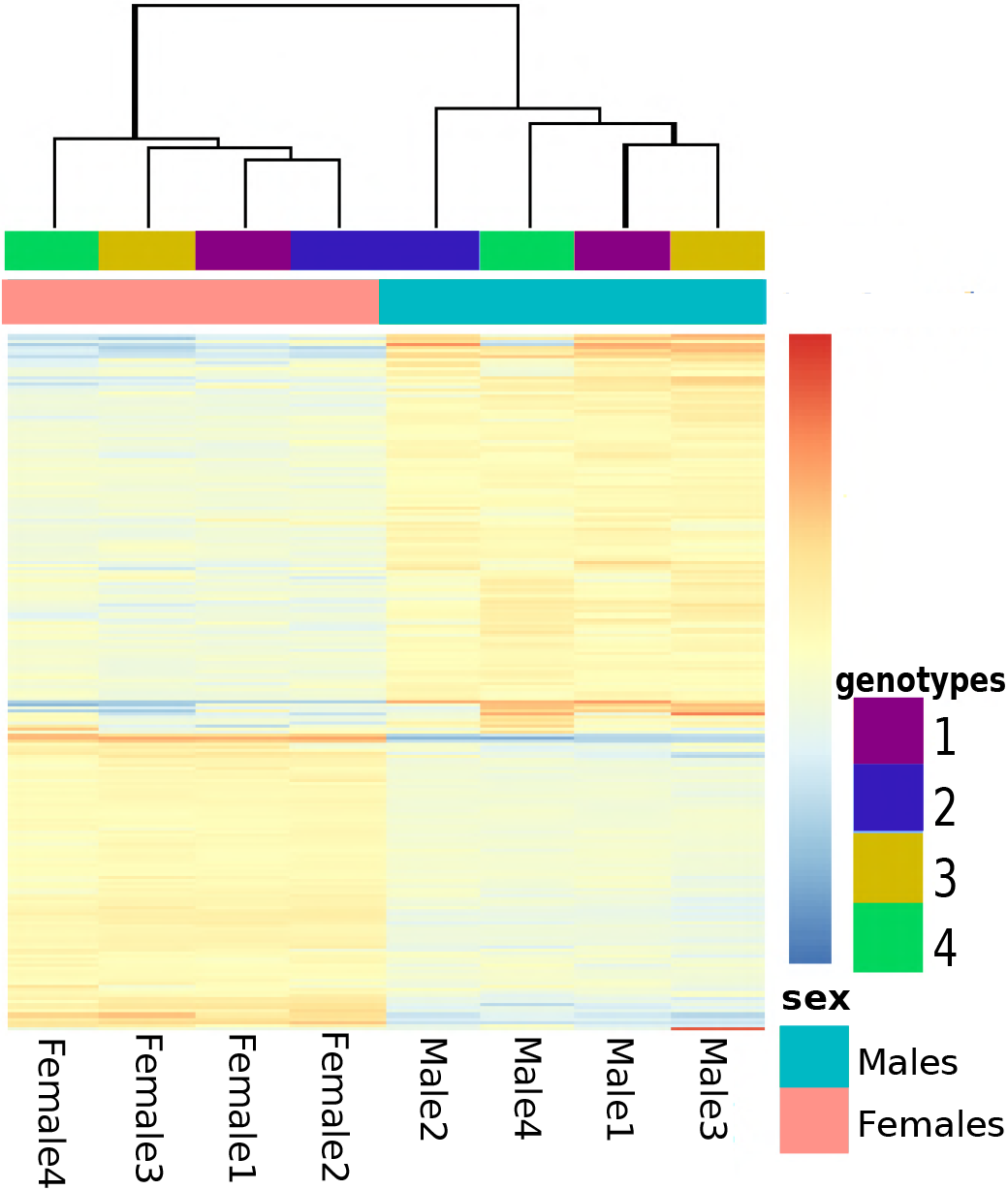
Heatmap showing the normalized expression levels of the sex-DE genes (p < 0.05) with at least a twofold expression difference between males and females. Each line represents a gene and each column specific sample (genotype and sex), with relative expression levels indicated by colour (from highly overexpressed, red to highly underexpressed, blue, as indicated by the scale to the right). The dendrogram above the sample columns indicates clustering according to the Euclidean distance matrix implemented in the pheatmap R

**Table 1:**
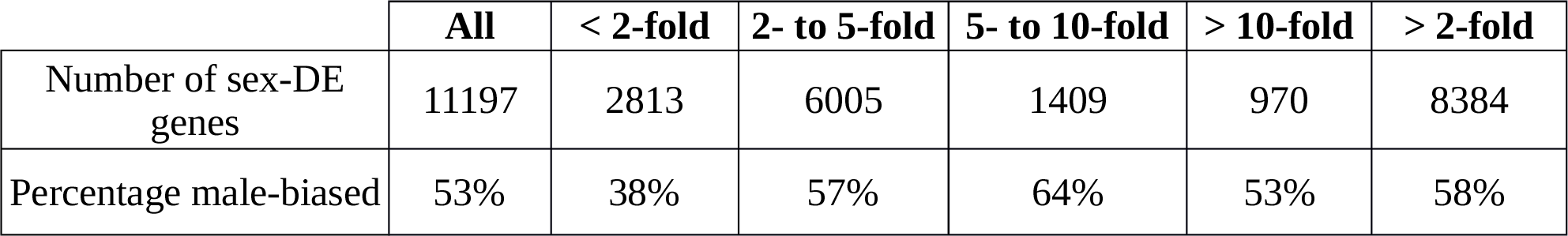
Numbers of significantly (adjusted p < 0.05) sex-DE genes in Daphnia magna, for different degrees of bias as well as percentage of genes with male-biased expression.

|  | All | < 2-fold | 2- to 5-fold | 5- to 10-fold | > 10-fold | > 2-fold |
| --- | --- | --- | --- | --- | --- | --- |
| Number of sex-DE genes | 11197 | 2813 | 6005 | 1409 | 970 | 8384 |
| Percentage male-biased | 53% | 38% | 57% | 64% | 53% | 58% |

### Comparisons of sex-biased gene expression among the three *Daphnia* species

The software OrthoFinder identified 7453 single copy, one-to-one orthologs present in all three species (**Table 2**). Among these, 5707 (76.5 %) were sex-DE in at least one species, and 707 genes (9.5 %) were sex-DE in all three species (**Fig. 2**, **Table2**). Only 32 of these 707 genes (4.5 %) showed a different direction of bias in one of the species. The remaining 675 genes were biased in the same direction in all three species, and we therefore refer to these genes as the core-set of sex-DE genes in *Daphnia*. Among the genes of the core-set, 75 % were female-biased (**Fig. 2**), and genes with a strong expression-bias between sexes were more likely to be included in this core-set (**Fig. 3**). Genes that showed significant sex-biased expression in only two out of the three species showed very similar patterns: a high proportion showed consistent bias (i.e., in the same direction in both species), and there was an excess of female-biased compared to male-biased genes (**Fig. 2**). The excess of female-biased genes was even observed among the genes with sex-biased expression in only one species (**Table S3**). This was not only the case in *D. pulex* and *D. galeata*, for which an excess of female-biased genes had been reported earlier (Huylmans *et al.* 2016), but also in *D. magna*, where this result contrasts with the slight excess of male-biased genes found when all 26’646 genes were considered (as opposed to only the 7453 genes, for which single-copy, one-to-one orthologs could be identified in the other two species). The list of the 7453 orthologs, as well as the data on sex-biased expression for the three species is given in the supplementary data (**Table S3**).

**Figure 2:**
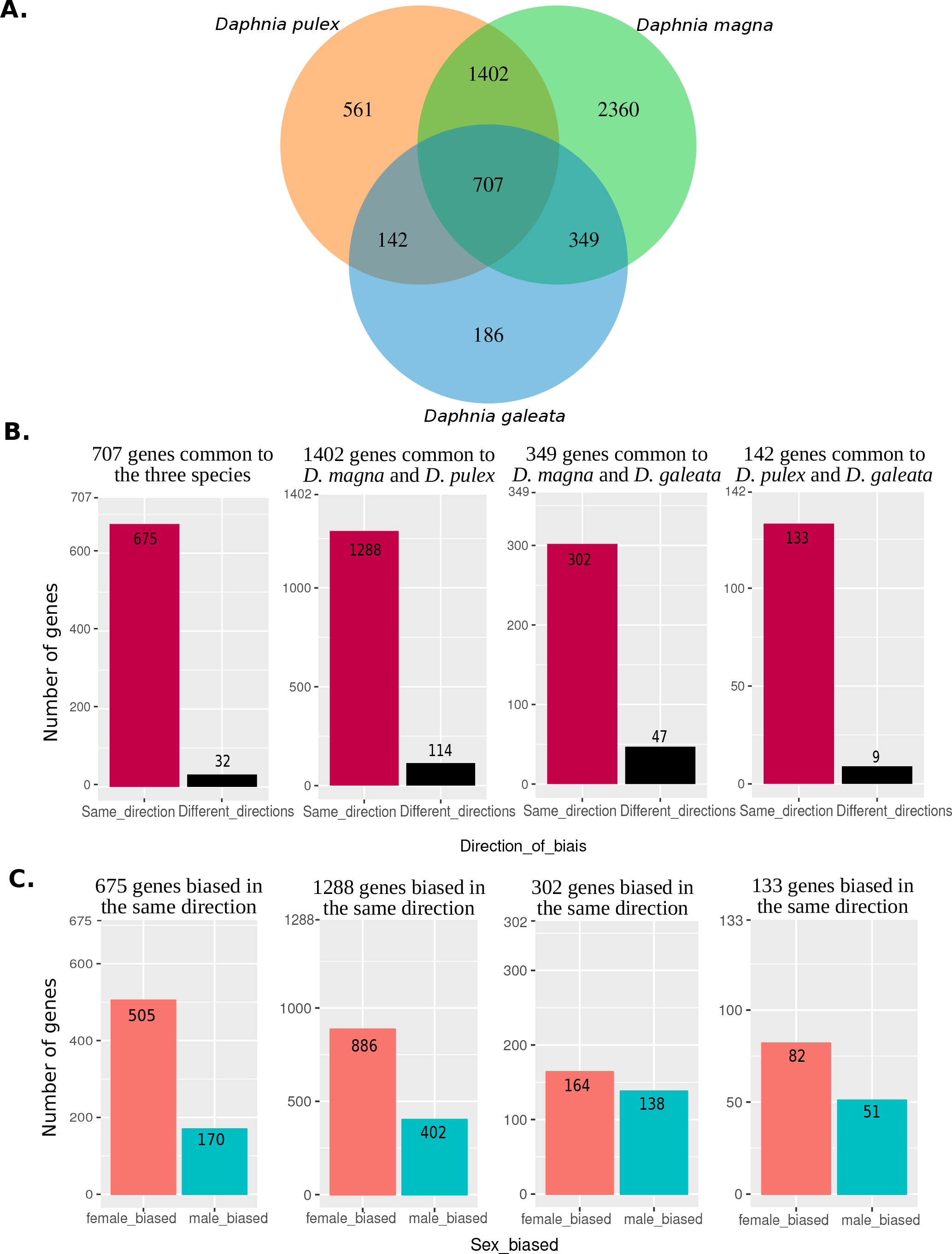
**A**. Venn diagram showing the number of sex-DE genes among the 7453 one-to one orthologs in each of the three species of *Daphnia*. **B.** Number of sex-DE genes being biased in the same vs. opposite directions. Panels from left to right: genes being sex-DE all three species (707 genes), genes being sex-DE only in *D. magna* and *D. pulex* (1402 genes), *D. magna* and *D. galeata*, (349 genes), and *D. pulex* and *D. galeata* (142 genes). **C.** Number of female-biased and male-biased genes in each of the four categories depicted in **Figure 2 B**. (only genes being biased in the same direction)

**Table 2:** Number of the one-to-one orthologous genes found by OrthoFinder in the three Daphnia species.

| Species | Number of protein sequences in the genome | Number of Sex-biased genes<sup>1</sup> | Proportion of sex-biased genes | Number of 1-to-1 orthologues | Number of sex-biased 1-to-1 orthologues |
| --- | --- | --- | --- | --- | --- |
| *D. magna* | 26646 | 11192 | 0.42 | 7453 | 4818 |
| *D. pulex* | 30940 | 6393 | 0.21 | 7453 | 2812 |
| *D. galeata* | 33555 | 5842 | 0.17 | 7453 | 1384 |

<sup>1</sup> as identified in the single species studies (current study, Colbourne et al 2011, Huylman

### Functional analysis of the core-set of sex-DE genes

Among the core-set of 675 orthologous genes that were consistently sex-DE in all three species, 592 had an annotated function. The major GO categories of these genes are shown in **Fig. 4**. This figure highlights that the largest fraction of genes belongs to the categories “cellular process”, “metabolic process”, “single organism process” and “biological regulation”. The results of the GO enrichment analysis are shown in **Fig. 5**. Enriched terms are linked to “RNA binding” processes, known to play a key role in post-transcriptional gene regulation (Glisovic *et al.* 2008; Cléry & Allain 2013) and, more generally, terms linked to “RNA”. Of the 592 genes only one gene has a GO term linked to sex determination or sex differentiation (which is neither more nor less than expected by chance): The gene “peptidyl-prolyl cis-trans isomerase FKBP4” has a female-biased expression in all three species and its GO term includes “male sex differentiation”. Moreover, among the same 592 core genes with a functional annotation, 14 were listed in the NCBI list of genes known to be involved in sex determination or differentiation pathways in other species (**Table S4**).

**Figure 3:**
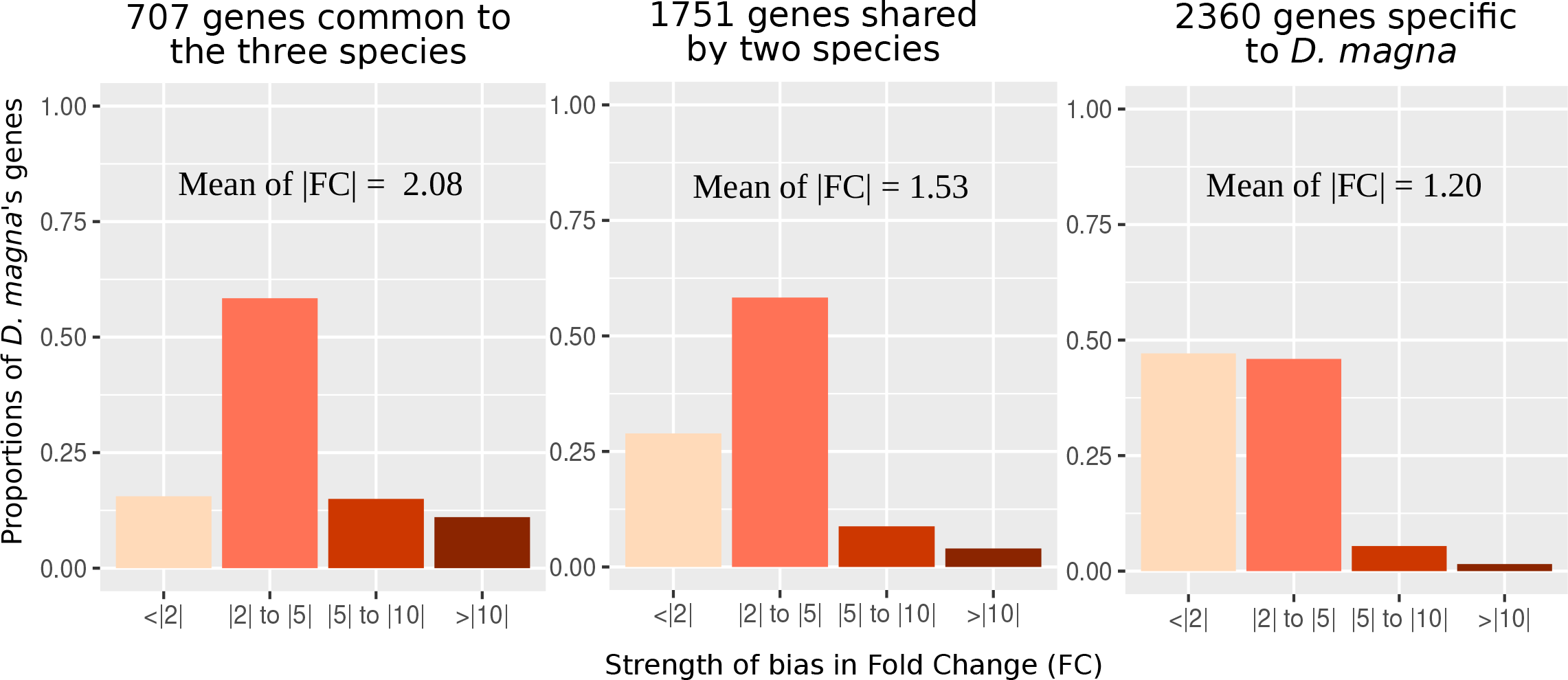
Proportion of genes with different degrees of sex-bias (the degree of sex-bias is summarized in four categories of fold change). Panels from left to right: genes being sex biased in all three species (707 genes), in two species (1751 genes), and only in *D. magna* (2360 genes). Only the 7453 genes, for which single-copy, one-to-one orthologs could be identified in all three species were considered for this analysis.

**Figure 4:**
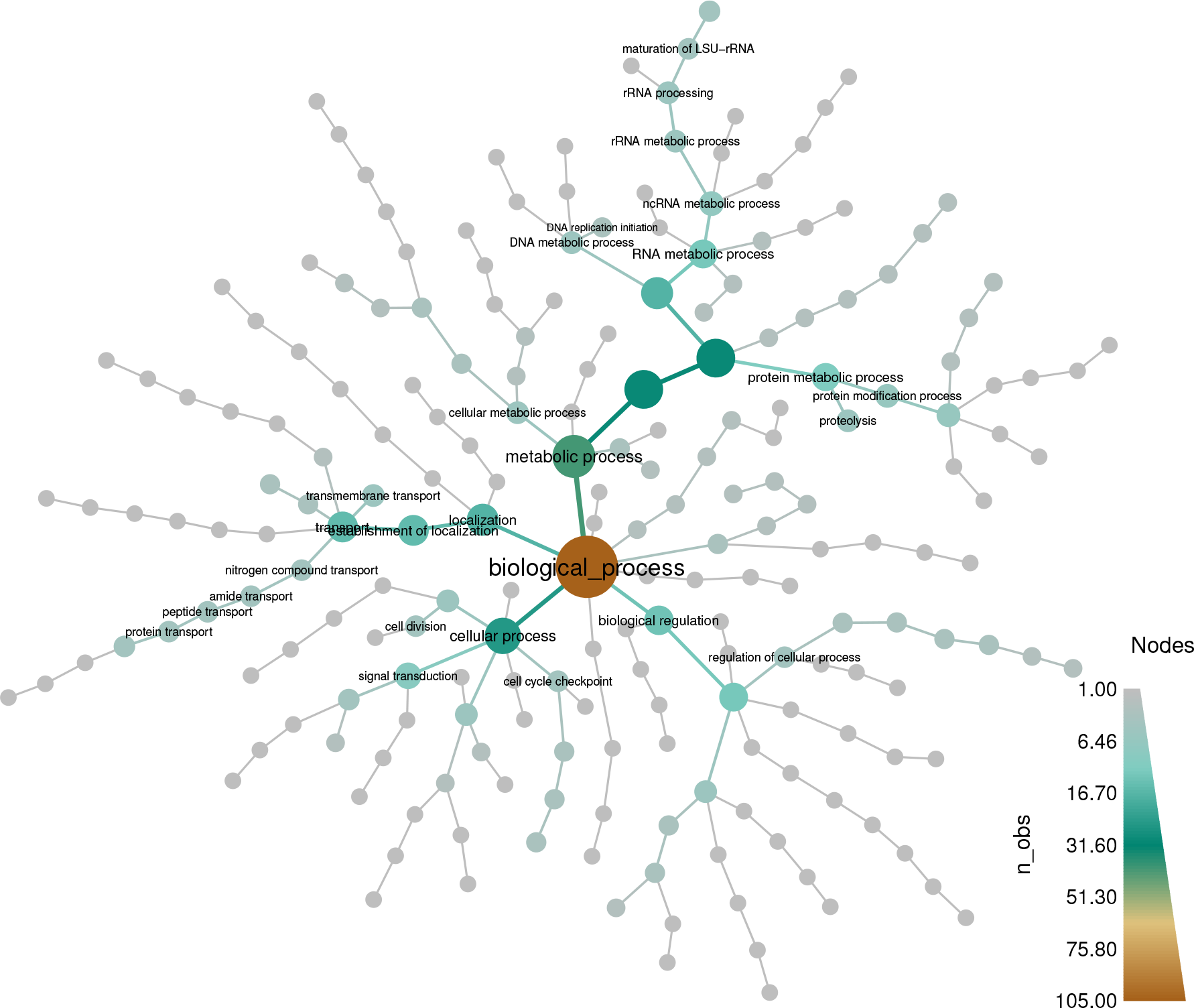
Composition and hierarchical organization of GO terms associated to the 675 genes with consistent sex-biased gene expression in all three species. n_obs: number of genes in the given GO category.

**Figure 5:**
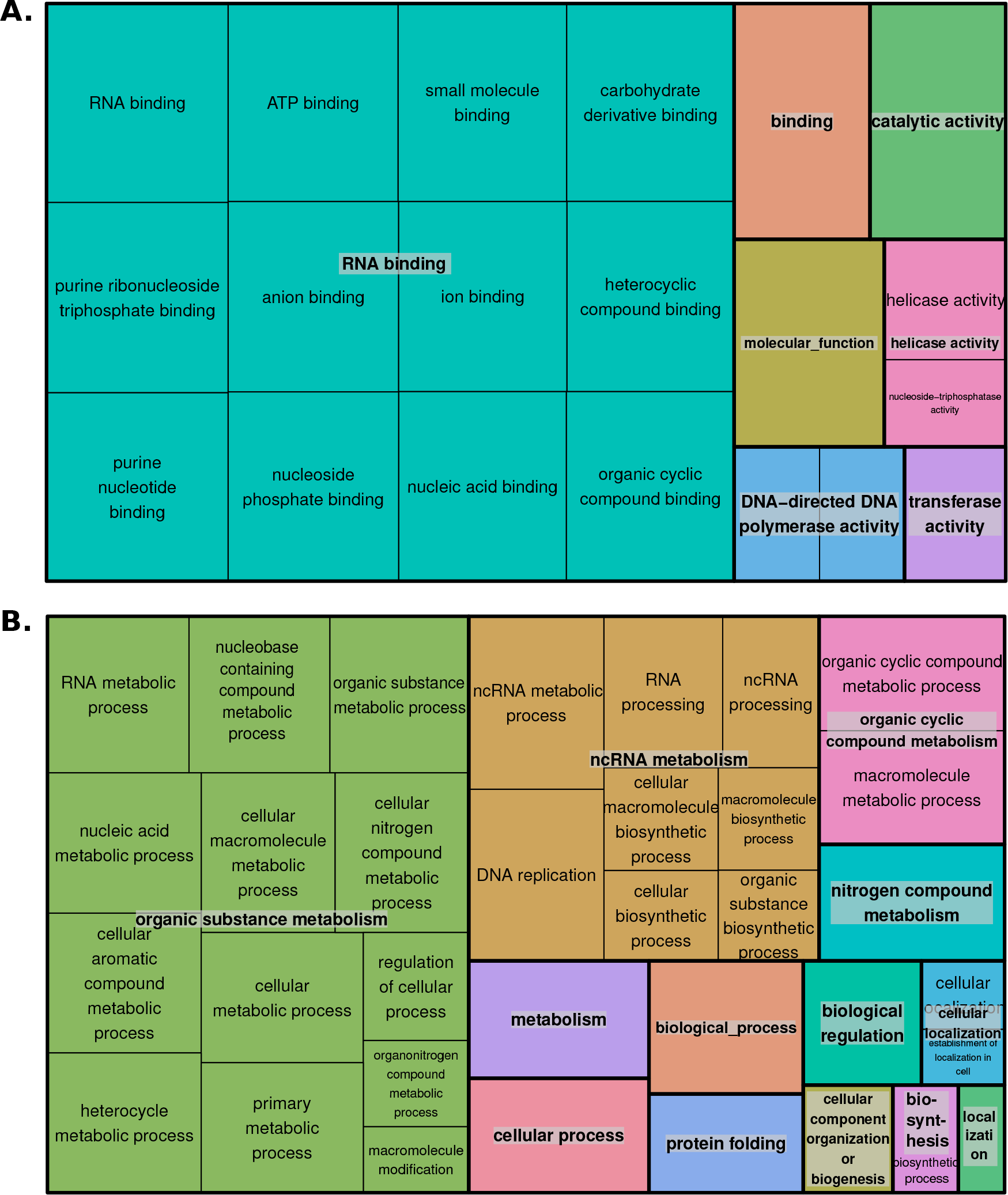
Enrichment analysis of GO terms among the core-set of genes compared to the entire list of *D. magna* genes. Shown are over-represented (Bonferroni-corrected *p* < 0.05) terms and functional categories (the latter distinguished by colour) for molecular functions (**A**) and biological processes (**B**). The size of each rectangle is proportional to the −log(*p*-value) for its category.

## Discussion

### Sex-biased gene expression in *Daphnia magna*

We found a very high number of genes being DE between males and females in *D. magna*. This result is largely congruent with the previous studies on other *Daphnia* species (Colbourne *et al.* 2011; Huylmans *et al.* 2016), but the overall number and proportion of genes that show sex-biased expression is higher in *D. magna* than in the other two species. Indeed, the proportion of genes with sex-biased expression in *D. magna* is similar to that reported for species with genetic sex determination (GSD). For instance, two early, microarray-based studies on *Drosophila melanogaster* found two-fold or greater expression differences between sexes in 30 % to 40 % of all genes (Parisi *et al.* 2004, Innocenti & Morrow 2010). A more recent study based on RNA-sequencing found that about two-third of genes showed sex-biased expression in multiple *Drosophila* species (genes with a less than two-fold change included). In *D. magna*, the proportion among all genes is 42 % (31 % with a greater than two-fold change). However, considering only the set of single-copy, orthologous genes (which likely contain a lower proportion of annotation errors, see below), the proportion is 65% (41% with a greater than two-fold change). Our results therefore suggest that these two systems, which are comparable in terms of body size and due to the fact that whole, adult animals were sampled, have similar proportions of sex-biased genes. More studies on comparable ESD-GSD species pairs will have to be investigated to determine the generality of this conclusion.

A recent study on *D. pulex* found that the annotation of the genome used here (Colbourne *et al.* 2011) likely contained a non-negligible fraction of pseudogenes or other false positives (Ye *et al.* 2017), and that the number of genes in the genome may be closer to 20’000 than the initially estimated ~31’000. It is possible that the current estimates of the total number of genes in the *D. magna* (~27’000) and *D. galeata* (~34’000) genomes may also be overestimates. Using these genomes as reference in a differential gene expression analysis may have affected the estimate of the proportion of differentially expressed genes only if the proportion of falsely annotated genes is different among the DE genes than among the non-DE genes. It is unclear if any such bias exists. However, two independent lines of evidence suggest that, if anything, such a bias has led to an underestimation of the proportion of sex-DE genes: First, the proportion of sex-DE genes was higher among the single-copy orthologs (which less likely contain annotation error) than among all genes in two out of three *Daphnia* species. Second, *D. pulex* and *D. galeata*, which both have higher estimated number of genes than *D. magna*, have a lower estimated proportion (~20 %) of sex-DE genes. Yet, the differences in the proportions of sex-DE genes among the three *Daphnia* species may also be explained by differences in methodology and statistical power. The *D. pulex* data (Colbourne *et al.* 2011) are based on a microarray study, a methodology known to be less sensitive for lowly-expressed genes than RNA-sequencing (Harrison, Wright & Mank 2012). The *D. galeata* study (Huylmans *et al.* 2016) was based on RNA-sequencing but used only two clonal lines as biological replicates. As mentioned by Huylmans et al (2016), this may have led to a rather low statistical power to detect sex-DE genes, especially among the considerable number of genes that showed expression differences between the two clones.

When comparing the proportion of genes with sex-biased expression with other studies, it is important to remember that we performed RNA-sequencing on whole animals and hence included all tissues present at that the time of sampling in adult males and females. Patterns of sex-specific gene expression are known to be tissue-specific in many cases (Ellegren & Parsch 2007; Toyota *et al.* 2017), with strongest differences being found in brains (at least in mammals) and, unsurprisingly, in gonad tissues (Mank 2009). Thus, it is difficult to compare our results with those in larger animals, where studies have mostly been carried out on specific tissues (e.g., 54.5 % of genes were found to be sex-DE in *Mus musculus* liver, (Yang *et al.* 2006)). Moreover, genes that do not show sex-biased expression may still differ in their expression patterns among tissues (Yang *et al.* 2006). Hence, when sampling whole animals, some of these genes may be identified as sex-biased because different tissues may occur in different proportions in males vs. females, for instance due to anatomical differences between sexes. It is possible that a part of the genes that were found to be sex-biased in our study and in other studies based on whole animals (e.g., *Drosophila*) are explained by such effects (i.e., sex-biased expression of these genes may be a consequence rather than a cause of the phenotypic differences between sexes (Mank 2017)).

Another factor that potentially contributes to an overestimation of the number of sex-DE genes is the fact that males were produced by artificially treating their mothers with the juvenile hormone analog MF (Huylmans *et al.* 2016). However, in a separate study we found that MF exposure changes expression levels of a much lower number of genes (only a few 100s) than were DE between sexes (Molinier *et al.*, in prep). Moreover, the males used in our experiment were exposed to MF only for three days when they were still oocytes inside the ovaries of their mothers up to one day after they were released from the brood pouch. It is thus highly unlikely that large proportions of the sex-DE genes are in fact explained by the effects of early MF exposure and would not have shown up in naturally produced males (which also involves exposure to a natural juvenile hormone produced by their mother).

Contrary to previous findings in *D. galeata* and *D. pulex* (which both show a clear excess of female-biased genes over male-biased genes), we observed a slight excess of male-biased genes in *D. magna.* It is currently difficult to say whether this difference between studies reflects a biological reality (i.e., difference between species within the genus) or whether it may be explained by some methodological differences between the studies. Interestingly, however, a strong excess of female-biased genes was recovered also in *D. magna* when investigating the subset of single-copy genes for which one-to-one orthologs could be identified in the other species. The excess of female-biased genes might be a general feature in the genus *Daphnia*, at least for single-copy genes that are sufficiently conserved for orthologs to be identified across the major sub-clades of the genus. On the other hand, the strong excess of female-biased single-copy one-to-one orthologs, suggests that the remaining genes show a substantial excess of male-biased expression, at least in *D. magna*. The remaining genes likely include many paralogs and other less conserved genes, for which the identification of orthologs is difficult. Hence, different evolutionary rates of genes with male-biased vs. female-biased expression could drive the observed patterns. Genes with male-biased expression evolve faster than female-biased genes in *D. pulex* and *D. galeata*, as well as in *Drosophila*, *Caenorhabditis*, and several mammal species (see reviews by Ellegren & Parsch 2007 and Parsch & Ellegren 2013). Faster evolution of genes with male-biased expression might be explained by positive selection being more common in these genes (Ellegren & Parsch 2007), and may lead to ortholog identification being more difficult in these genes, which may explain or at least contribute to the observed difference between the two sets of genes in *D. magna.*

### Comparisons of sex-biased gene expression among the three *Daphnia* species

The 7453 single-copy, one-to-one orthologs present in all three species represent less than 30 % of all genes used in the *D. magna* analysis. However, as pointed out above, the number of genes predicted by the current genome annotation used in this study may in fact be a rather strong overestimation of the true number of genes present in the species. Secondly, the software identified a considerably higher number of orthogroups, which, however, also include paralogs. We decided to restrict the analysis to single-copy, one-to-one orthologs because interpretation of expression patterns in paralogs is much less straightforward. For instance, if a sex-DE gene is single-copy in one species but has two paralogs in another species due to duplication after speciation, it is difficult to say which one of the two genes is more homologous in function, and hence should also be sex-DE if the gene belongs to the core-set of genes with sex-biased expression in the genus (Koonin 2005).

Only a low percentage of the genes found to be sex-biased in two and especially in three species were biased in different directions. Moreover, genes with a higher expression bias were more likely to be sex-DE in two or three species than just in one. While the latter observation may in part be explained by issues of statistical power (genes with a low degree of sex-bias have a lower probability to be detected), both observations nonetheless suggest that the 675 orthologous genes that were found to be consistently sex-DE in all three species indeed represent a robust core-set of sex-biased genes in the genus *Daphnia.* It is likely that some genes that were DE between a pair of species but not in all species should also have been included in this core-set, as differential expression may have been non-significant in one species just due to a lack of statistical power. Indeed, the three studies differ in methodology (microarray vs. RNA-Sequencing), number of biological replicates, aspects of data analysis, etc (see above). In addition, the quality of the genome assemblies and annotations used to analyze these data may also differ between species. These differences may also explain some of the between-species differences in the number and proportion of sex-DE genes.

### Functional analysis of the core-set of sex-DE genes

Our study identified a core-set of genes for which sex-biased expression is probably conserved in the genus *Daphnia*. Hence these genes may play a fundamental role in determining and maintaining male vs. female phenotypes in this genus with environmental sex determination. The functional analysis of these genes identified the gene “peptidyl-prolyl cis-trans isomerase FKBP4” with GO term “male sex differentiation”, an immunophilin protein with peptidylprolyl isomerase and co-chaperone activities. It is a component of steroid receptors heterocomplexes and may play a role in the intracellular trafficking of hetero-oligomeric forms of steroid hormone receptors between cytoplasm and nuclear compartments. Steroid receptors initiate the signal transduction for steroid hormones, including sexual hormones such as oestrogen and androgen (e.g., Voigt *et al.* 2009). Their role in sex dimorphism, also in species with environmental sex determination, thus makes sense, though the role of this particular gene has not yet been investigated in *Daphnia*. The 14 additional genes whose functional annotation matched of the descriptors of the genes on the NCBI list of genes involved in sex determination/differentiation pathways may represent further fundamental genes involved in sex determination or sex differentiation in *Daphnia*. They contain functions known to be implicated in sexual development in other species, such as the “Beta-catenin 1” which is a key transcriptional regulator of the canonical Wnt-signaling pathway, known to be implicated in female reproductive development in mammals (Bernard & Harley 2007, Liu, Bingham & Parker 2008). We also found the “fibroblast growth factor receptor”, receptor of Fgfs (fibroblast growth factors), whose function may be involved in sex determination and reproductive system development in many species and appears to be highly conserved (Colvin *et al.* 2001). Finally, the gene ‘ovarian tumor” is also known to be involved and required in the determination of the sexual identity of female germ cells (Pauli, Oliver & Mahowald 1993). Our results suggest that these genes are involved in maintaining phenotypic differences between sexes, also at the adult stage, at last in *Daphnia.* A large proportion of the identified core genes have unknown function. Therefore, we need more functional annotations, especially also on more closely related species, as well as sex-specific expression data on earlier developmental stages before we can obtain a clear mechanistic picture of sex determination and sex differentiation in *Daphnia* (Kato *et al.* 2011).

In conclusion, our study provides data on sex-biased gene expression for the model organism *D. magna* and for *Daphnia* in general, specifically by identifying a core-set of sex-DE genes in all three major subclades of the genus. More generally, our results suggest that the proportion of genes with sex-biased expression in ESD species is not lower than in species with GSD.

## Acknowledgements

We thank the Zoo of Moscow and N. I. Skuratov for sampling permits, and David Frey for help with maintenance of indoors cultures. We thank the Department of Biosystem Science and Engineering of the ETH Zurich, in particular C. Beisel and I. Nissen for Illumina sequencing, and we gratefully acknowledge support by M-P. Dubois, the platform Service des Marqueurs Génétiques en Ecologie at CEFE, and the genotyping and sequencing facilities of the Institut des Sciences de l’Evolution-Montpellier and the Labex Centre Méditerranéen Environnement Biodiversité (CeMEB). We thank Mathilde Cordellier and John Colbourne for sharing data and information and two anonymous reviewers for their comments. We thank the University of Fribourg, the Montpellier Bioinformatic Biodiversity plateform and the Labex CeMEB for access to high-performance computing clusters. The sequence data for the *D. magna* genome project V2.4 were produced by The Center for Genomics and Bioinformatics at Indiana University and distributed via wFleaBase in collaboration with the *Daphnia* Genomics Consortium (project supported in part by NIH award 5R24GM078274-02 *“Daphnia* Functional Genomics Resources”). Our work was supported by the Swiss National Science Foundation (Grant no. 31003A_138203), the Russian Foundation of Basic Research, and the European Union (Marie Curie Career Integration Grant PCIG13-GA-2013-618961, DamaNMP). YG was supported by RFBR grant 16-04-01579.

